# Ethanol-assisted core-shell microparticles for enzyme stabilization with precise size control

**DOI:** 10.64898/2026.05.05.722948

**Authors:** Eunhye Yang, Waritsara Khongkomolsakul, Younas Dadmohammadi, Alireza Abbaspourrad

**Author notes:** Corresponding authors, Alireza Abbaspourrad.

## Abstract

In vegetarian diets, phytate is known to disrupt the adsorption of minerals. Fortifying foods with phytase, a therapeutic enzyme known to mitigate phytate, might increase the uptake of important nutrients. Phytase is susceptible to environmental stress such as heat and acidic conditions encountered during food processing. Therefore, we developed and optimized a core-shell microparticle composed of a phytase-chitosan core and a shell consisting of cross-linked alginate-κ-carrageenan. Ethanol was used to precipitate the microparticles, and the ethanol concentration was optimized along with the chitosan and phytase ratio and the alginate-carrageenan concentration, to form stable core-shell microparticles. The optimized core-shell microparticles have a loading capacity of 32.7% with a high encapsulation efficiency of 80.3% and uniform micro-size with a diameter of 3.2 µm and a poly-dispersity index of 0.178. Loaded phytase retained 62.7% enzymatic activity after heat treatment and digestion conditions. These results indicate that core-shell microparticles are suitable for retaining enzyme activity within the food matrix under typical food processing conditions.

**Highlights:**

- Development of size-controlled core-shell microparticles to protect phytase
- Phytase-chitosan microparticles are surrounded by an alginate-κ-carrageenan shell
- Optimization achieved 32.7% loading capacity with a uniform size of 3.2 µm
- Core-shell microparticles retained 62.7% enzyme activity after heat and digestion
- Phytase powder (2 mg) is required for a single maize meal

## 1. Introduction

Therapeutic enzymes, including phytase, lipase, and protease, serve therapeutic purposes, such as diabetes control, anti-inflammation, and anti-obesity (Tandon et al., 2021). A recent survey indicated that consumers prefer drugs and supplements that can be administered orally (Alqahtani et al., 2021). The therapeutic enzymes are difficult to administer orally because they are heat sensitive, can be destroyed by digestive enzymes, and are unstable in low pH conditions. We selected phytase, a therapeutic enzyme used to prevent mineral deficiency by hydrolyzing phytate and releasing calcium, phosphorus, and other nutrients, as a model enzyme to be developed for oral administration. To do this, we will need to encapsulate phytase such that it can make it through the digestive tract to the small intestine, where it can be absorbed and used.

Polysaccharides interact with enzymes through electrostatic forces, and the interactions can be modulated by pH and ionic strength. Polysaccharides have been shown to protect an enzyme’s structural integrity from external environmental conditions, thus retaining enzyme functionality (Bialas et al., 2021; Weng et al., 2023). In addition, polysaccharide-based enzyme encapsulation allows for a controlled release triggered by pH changes and can be developed to be responsive to the specific target site of interest. This approach has several advantages, including that it is consumer-friendly (natural polymers) and biocompatible.

Recent studies on enzyme encapsulation using polysaccharides, including hydrogels, nanoparticles, and microspheres, are driven by different mechanisms such as electrostatic assembly (Weng et al., 2023), coacervation (Li et al., 2022; Liu et al., 2023), and spray drying (J. Wang et al., 2023). The enzyme-polysaccharide complex produced using such methods consistently suffers from reduced enzyme activity despite having a protective effect against digestion conditions (Song et al., 2022). Unlike other proteins, enzymes have specific functionality only when the active site is entirely open, so it is perhaps not unexpected that the decreased activity is from the active site being blocked by the polysaccharide or at least the geometry of the site tweaked.

Previously (Yang et al., 2024), we reported a polysaccharide-based core-shell hydrogel bead that protected the enzyme from the digestive tract, but also maintained enzyme activity by reducing the direct interactions between phytase and the polysaccharide.

Despite being able to protect and maintain function in our previous attempts at core-shell hydrogel beads, the method resulted in a rather large-sized bead (minimum diameter ∼2-3 mm), and the method was not suitable for running at scale. For food applications specifically, a micro-sized bead is essential. Therefore, we developed a new strategy to manufacture enzyme-polysaccharide-based microscale particles while maintaining the core-shell structure based on anti-solvent precipitation. We hypothesized that a positively charged enzyme-polysaccharide complexes can serve as the core and interact with an anionic polymer to form core-shell particles through electrostatic interactions. Anti-solvent precipitation was used to facilitate particle formation and improve size uniformity.

Antisolvent precipitation is used to produce nano- and micro-encapsulated particles by reducing the solubility of one substance in a solvent system (Zhang et al., 2022). This method can produce particles on the nano- to micrometer scale with large surface area and with the flexibility to control particle size and distribution. Particles on this scale are likely to affect the release behavior of the encapsulated compounds and their physicochemical stability. In addition, it can be used for heat-sensitive compounds because it can help particle formation under mild conditions without high-temperature processing. Therefore, it is a promising strategy for enzyme encapsulation, maintaining enzymatic activity. Recent studies (Caicedo Chacon et al., 2023; Liu et al., 2024) highlight its effectiveness in various applications, including pharmaceuticals and industrial processes. The choice of antisolvent and process conditions is a critical factor. Ethanol is the most common organic solvent for antisolvent precipitation. Ethanol-based protein aggregation impacts solubility and facilitates the formation of nano- and microparticles. While ethanol is commonly used, it is important that the ethanol concentration not be too high, as it may permanently damage the enzyme structure.

We report the preparation of core-shell microparticles such that the core is a phytase-chitosan microparticle using ethanol anti-solvent precipitation. and the shell is composed of anionic polysaccharides. The protective effect of the microparticle was measured by the retention of enzyme activity. The particle size was on the microscale. The preparation and optimization of core-shell microparticles with phytase, using ethanol as an anti-solvent precipitation controlled the particle size. The core-shell structure protected phytase from environmental stressors such as heat and acidic conditions, and it was able to hydrolyze phytate in a real food matrix.

## 2. Materials and methods

### 2.1. Materials

Potassium chloride (KCl, > 99 %), alginic acid sodium salt from brown algae (medium viscosity), sodium hydroxide anhydrous (NaOH, > 97 %), calcium chloride dihydrate (CaCl_2_, > 99.5 %), ammonium molybdate tetrahydrate, trichloroacetic acid (TCA), and L-ascorbic acid were purchased from Sigma-Aldrich (St. Louis, MO, US). Phytase from *Aspergillus niger* was supplied by DSM company (Heerlen, Netherlands). Pierce^TM^ rapid gold BCA (bicinchoninic acid) protein assay kits were purchased from Thermo Fisher Scientific (Liverpool, NY, US). Sodium phytate (95 %) was purchased from Astatech Inc. (Bristol, PA, US). κ-Carrageenan was purchased from TIC GUMS (Westchester, IL, US). Chitosan from *Aspergillus niger* was purchased from Sarchem Laboratories (Farmingdale, NJ, US). Sulfuric acid (95-98 %) was purchased from Fluka Honeywell (Charlotte, NC, US).

### 2.2. Preparation of core-shell microparticles

The alginate/κ-carrageenan stock solution (1:1 (w/w)) was incubated at 60 °C overnight. Chitosan and phytase stock solutions were prepared separately. The two solutions (chitosan and phytase) were first mixed at room temperature. Calcium chloride and potassium chloride were also dissolved separately and then added as cross-linker solutions to make each final concentration (chitosan 4 w/v%, phytase 6 w/v% or 8 w/v%, calcium chloride 0.2 M, and potassium chloride 0.05M) at room temperature (∼25 °C). After combining chitosan and phytase/linker solutions, ethanol was subsequently added at various ratios. The resulting phytase-chitosan microparticle-containing solution was then homogenized (probe-type, Ultraturrax) for 5 min at 12,000 rpm. The core-shell microparticle was then produced by mixing the microparticle solution and alginate-κ-carrageenan solution (∼0.15-0.45 v/v%) in a 1:2 ratio. The phytase-chitosan core-shell microparticle solution that was produced was again homogenized for 5 min at 12,000 rpm. The final core-shell microparticle solution was spray-dried (LabPlant SD-Basic, spray dryer (North Yorkshire, U.K.)) to produce a powder sample for further analysis (SEM) and to maintain stability during storage. Solutions were fed into the drying chamber through a peristaltic pump with an inlet air temperature of 170 °C and outlet air temperature of 70 °C with an airflow of 600 L h^−1^. The feeding rate was 16.7 mL min^−1^, with an atomization pressure of 20 psi. The powder-formed phytase-loaded core-shell microparticle was then reconstituted in distilled water at a protein concentration of 0.1∼1.0 w/v%.

### 2.3. Encapsulation efficiency, released phytase, and loading capacity

We used a Pierce rapid gold BCA protein assay kit to determine the amount of phytase (Steć et al., 2023). The sample solution (20 μL), containing the phytase, was treated with a 200 μL mixture of cupric sulfate and a copper chelator in a microplate well and incubated for 5 min at room temperature (∼25 °C). The absorbance of the treated supernatant was measured at 480 nm using a plate reader (SpectraMax iD3, Molecular Devices, CA, US) to determine the encapsulation efficiency, released amount, and loading capacity. Encapsulation efficiency and loading capacity of phytase were evaluated by isolating the free phytase by centrifugation at 6,700*g* for 10 min after preparation or incubation. The encapsulation efficiency was calculated as the following equation (**Eq.1**):

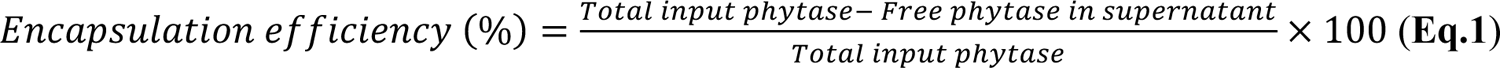

The loading capacity was calculated from the dehydrated mass and the loaded amount of phytase as the following equation (**Eq.2**):

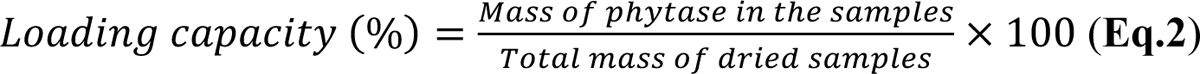

### 2.4. Phytase activity assay

A modified version of a previously reported method (Duru Kamaci & Peksel, 2021) was used to assess the phytase activity in the microparticles. The 150 μL substrate solution containing 0.44 mM sodium phytate in 100 mM sodium acetate buffer (pH 5.0) was pre-heated for 10 min in a 50 °C water bath. A 150 μL aliquot of either the free phytase solution or the phytase-loaded core-shell microparticle suspension was added to the substrate. The enzymatic reaction occurred for 30 min at 50 °C and was stopped by the denaturing agent (400 μL of 15 w/v% trichloroacetic acid (TCA) solution). After cooling down and centrifugation (6,700*g* for 10 min), 100 μL of the resulting supernatant was put in the mixture of 1 mL of the color reagent (1:1:3 (v/v/v) composed of 10 w/v% ascorbic acid, 2.5 w/v% ammonium molybdate, 1 M sulfuric acid solution) and 900 μL of distilled water. This mixture was incubated for 15 min at 50 °C before cooling to room temperature for 10 min. The absorbance of the mixture was measured at 820 nm using a UV–Vis spectrophotometer (UV-2600, Shimadzu, Kyoto, JP). We used a standard curve based on phytase solution to determine the residual activity and thermal stability. In the case of thermal stability, the residual activity was measured after heating the free phytase or loaded phytase solution for 12 min at 100 °C.

### 2.5. Food matrix and digestion condition

#### 2.5.1. In vitro digestion model

Based on the INFOGEST digestion protocol, we prepared a gastric buffer using a pH 2 KCl-HCl buffer with 1 mg/mL pepsin. To simulate gastric digestion, we mixed the sample solution and gastric buffer in a 1:1 (v/v) ratio and incubated the mixture at 37 °C for 2 h.

#### 2.5.2. Maize porridge

We prepared a porridge-like corn starch solution based on the following general recipe: 20 g corn starch was mixed with 30 mL water, and 166 mg sodium phytate was added as the substrate. The solution was stirred at 100 °C and 500 rpm for 10 min. We add a phytase-loaded core-shell microparticle powder to the simulated porridge, stirring at 60 °C and 500 rpm for 10 min. Then, this mixture was mixed with gastric buffer in a 1:1 ratio, stirring at 37 °C for 60 min. After incubation, 1 mL of the sample solution was collected and centrifuged (6,700*g* for 10 min). The supernatant was collected, and the phosphate concentration was measured following the same procedure used in the phytase activity assay, including the color reagent addition and absorbance measurement at 820 nm.

### 2.6. Physicochemical characterization of phytase-loaded microparticles

#### 2.6.1. Fourier transform infrared spectroscopy (FTIR)

Intermolecular interactions between phytase, chitosan, κ-carrageenan, and alginate were investigated using FTIR spectroscopy. FTIR analysis was performed on an IRAffinity-1S Spectrometer (Shimadzu Corporation, JP) fitted with a single-reflection attenuated total reflectance (ATR) accessory. The measurement was conducted with an average of 32 scans from 400 to 2000 cm^-1^ (resolution = 4 cm^-1^) after testing a background without a sample.

#### 2.6.2. Size distribution, polydispersity index, and particle size measurement

Particle size and polydispersity index of phytase-chitosan complex, microparticles, and core-shell microparticles will be measured using dynamic light scattering and particle electrophoresis on a Zetasizer Nano-ZS (Malvern, United Kingdom).

#### 2.6.3. Scanning electron microscope (SEM)

The size and uniformity of core-shell microparticles were observed using a Zeiss Gemini 500 scanning electron microscope (Zeiss, DE). After spray-drying, dehydrated microparticles were positioned on a carbon-taped stub and coated with gold using a sputter coater (Denton Desk V, NJ, US). The resulting samples were scanned and imaged using a secondary electron detector with a 20.0 μm aperture (accelerating voltage = 1 kV).

### 2.7. Statistical analysis

The data are presented as means and standard deviations of triplicates and analyzed using Analysis of Variance (ANOVA). The Tukey HSD comparison test evaluated the differences between mean values (p < 0.05). All statistical analyses were performed using JMP Pro 17 (SAS Institute, US) and plotted by GraphPad Prism 10 (GraphPad Software Inc., US).

## 3. Results and Discussion

### 3.1. Anti-solvent precipitation with ethanol

The required concentration of ethanol to precipitate microparticles of chitosan, phytase, and phytase-chitosan complex was investigated. Our criteria for the appropriate concentration were the formation of microparticles with uniform size and a positive charge so that they could bind with alginate and κ-carrageenan to form the core-shell structure in the second step. We considered three aspects of ethanol concentration: impact on chitosan gelation, protein aggregation, and protein denaturing.

The impact of ethanol on chitosan showed that in high ethanol content chitosan gels, due to cross-linking between the cross-linker agent that is also present (Ca^2+^) and chitosan (Pantić et al., 2023). The viscosity and turbidity of the chitosan solution increased when the ethanol concentration was above around ethanol 40 v/v%. Gelation of the chitosan inhibits the interaction with the anionic polymers, like the alginate/κ-carrageenan in the secondary shell process. Therefore, the ethanol concentration must be lower than the chitosan gelation point.

The phytase-chitosan complex has a bigger particle size than a single enzyme; it can become a micro-sized particle even if only 2-3 complexes are gathered. Therefore, optimal phytase-chitosan microparticles are formed at lower ethanol concentrations than those for raw phytase.

Protein aggregation is induced by adding ethanol, although it varies depending on the protein type, size, and stability. Some proteins are permanently denatured by high ethanol concentration, causing the structure and function to be lost. Therefore, identifying the ethanol concentration range that induces aggregation to form the desired size microparticles while maintaining structural stability is necessary. Protein aggregation and overall increase in particle size were evaluated by measuring absorbance at 500 nm, which reflects increased light scattering associated with larger aggregates..

The chitosan solution was transparent up to 30 v/v% ethanol concentration, then at 40 v/v% the solution showed an increase in turbidity and increased viscosity indicative of gelation (**Figure 1(a)**). The phytase-chitosan complex showed increased turbidity across all concentrations of ethanol starting at 10 v/v% ethanol (**Figure 1(b)**). Raw phytase’s turbidity started to increase at ethanol concentrations of 25 v/v% (**Figure 1(c)**). The turbidity of the complex plateaued at 30 v/v% ethanol, but the ethanol concentration range can be divided into three regions based on the slope: (1) 5-15 v/v%, (2) 15-25 v/v%, and (3) 25-40 v/v%. Region (1) is attributed to the initial association of intact enzyme molecules. Region (2) represents a translation or mixed state where both intact enzyme associations and denatured enzyme complexes coexist. Region (3) corresponds to the complexation of denatured enzymes, either with each other or with remaining intact enzymes

**Figure 1.**
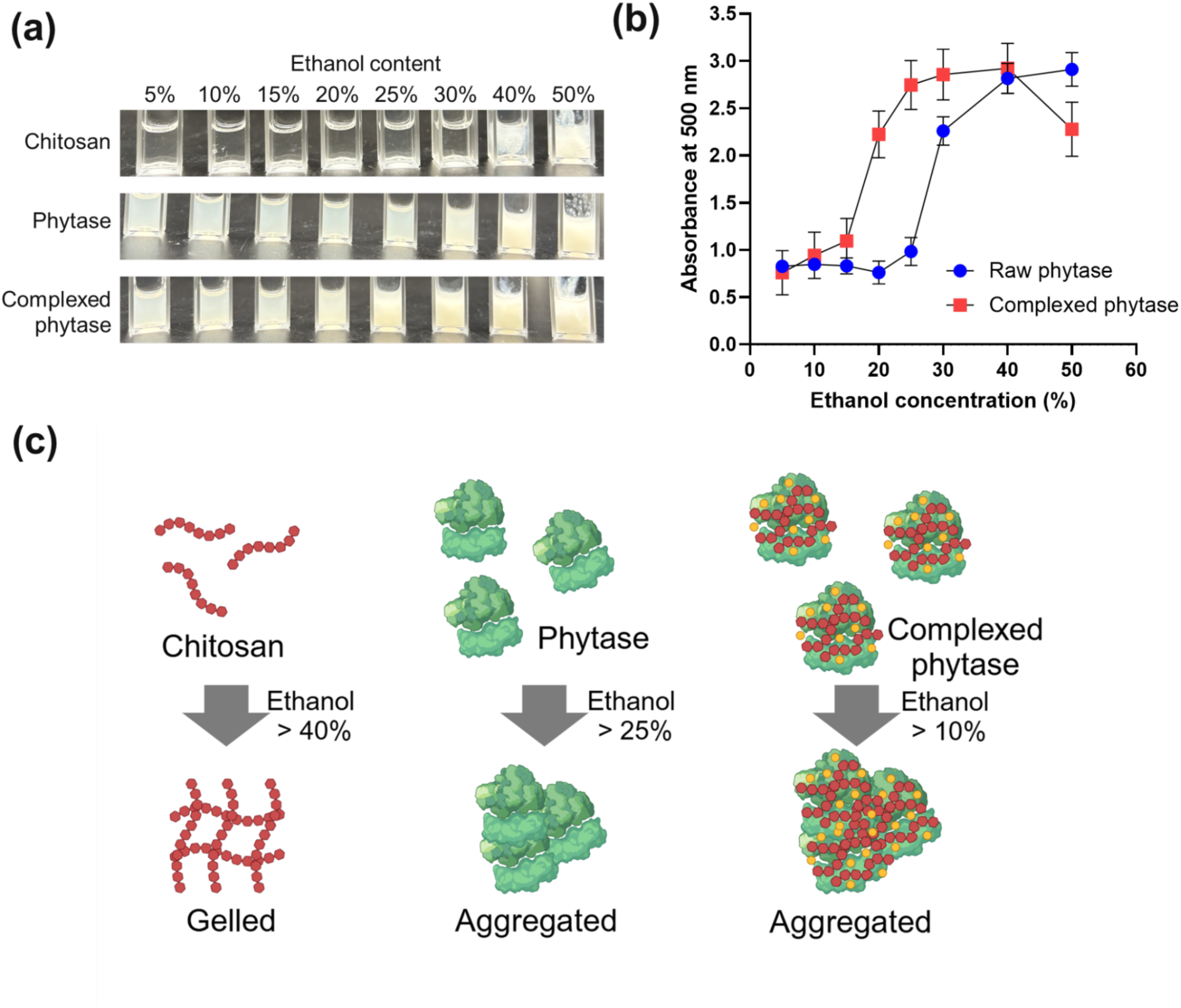
Effect of ethanol concentration on chitosan, phytase, and chitosan-phytase complex. (a) Appearance of chitosan, phytase, and chitosan-phytase complex solution depending on the ethanol concentration. (b) Turbidity of phytase and chitosan-phytase complex solution depended on the ethanol concentration. (c) Graphical illustration representing the gelation/precipitation appearance of chitosan, phytase, and chitosan-phytase complex.

Samples with ethanol concentrations of 10, 20, and 30 v/v% were selected to represent each region. Turbidity was compared under three dilution conditions designed to distinguish the effects of ethanol concentration and dilution: (1) dilution while maintaining a high ethanol environment, (2) dilution with water to reduce ethanol content, and (3) dilution to an intermediate ethanol concentration. In all cases, the total volume and complex concentration were kept constant. If the turbidity of (b) is not decreased as much as the turbidity of (c), the agglomerated particles are the result of the complexation of denatured protein, and it can be interpreted that the high concentration of ethanol permanently damages the structural stability and functionality of the phytase. At ethanol concentrations between 25-40 v/v%, phytase was denatured, and agglomeration did not drop, so turbidity did not decrease ( **Figure 2(a)**). In the 15-25% v/v% range, a partial reduction in turbidity upon dilution suggests that some of the aggregates are reversible. In contrast, the 5-15% v/v% range showed a more substantive decrease in turbidity, indicating that aggregation was largely reversible. The better preserved structural integrity of phytase was found at lower ethanol concentrations, therefore, ethanol concentrations of 15 v/v% or lower are preferable.

**Figure 2.**
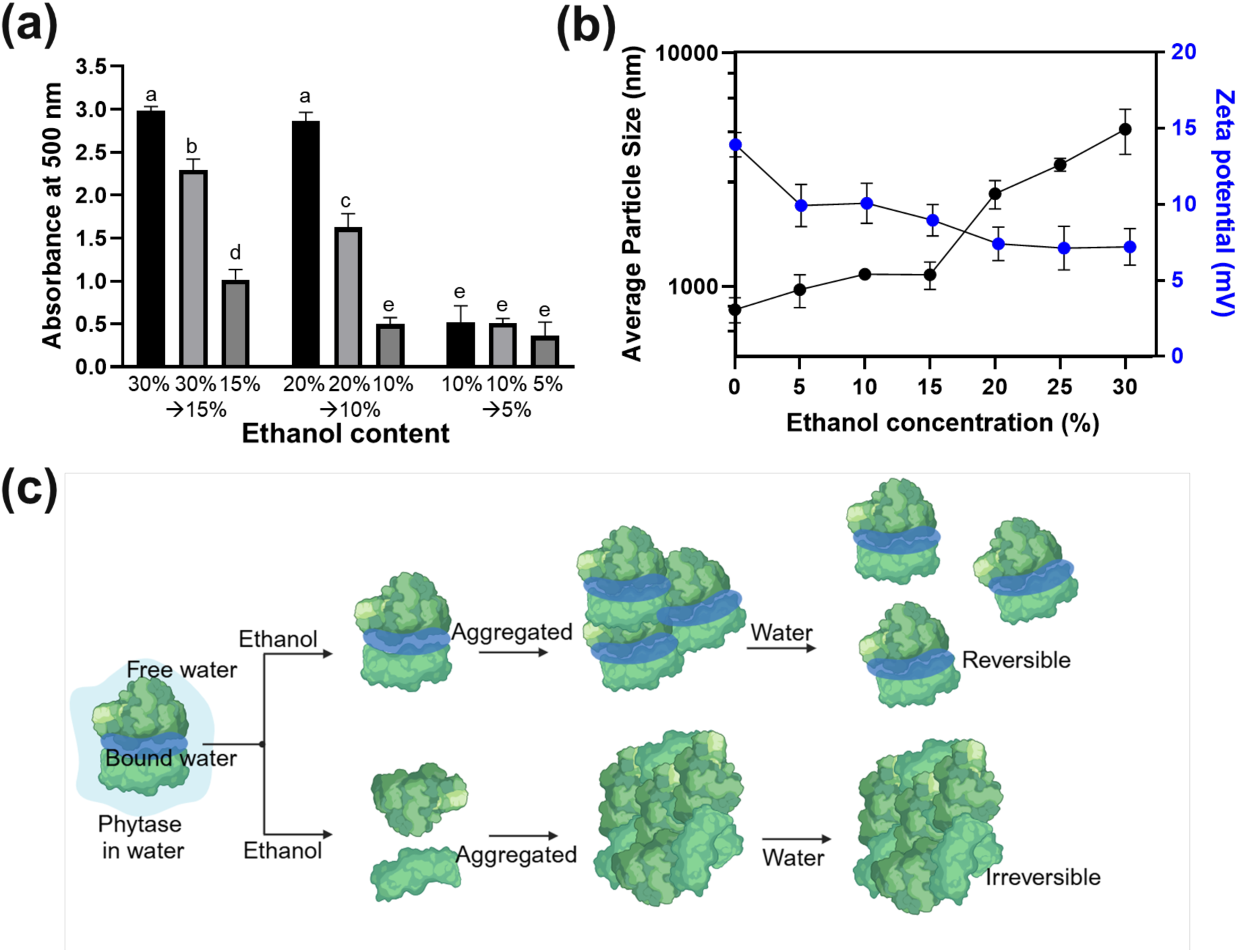
Effect of ethanol concentration on chitosan-phytase microparticles. (a) Turbidity of chitosan-phytase complex solution depending on the ethanol concentration and dilution conditions. A connecting letter plot was used at the significance difference level of p ≤ 0.05. (b) Graphical illustration representing the effect of ethanol on the free/bound water around the phytase. (c) Average diameter and zeta potential of chitosan-phytase microparticles depending on the ethanol concentration.

Finally, the particle size and charge when the ethanol concentration is within this range, 5-15 v/v%, confirmed that the complex would react with the anionic polymers in the second process, and the particle size was in the micrometer range. At an ethanol concentration of 15 v/v%, the particle size was larger than 1 µm (z-average diameter: 1.2 µm), which was consistent with the turbidity results. Also, the zeta potential of the microparticle with ethanol 15 v/v% was +9.6 mV (**Figure 2(b)**). To maintain electrostatic repulsion between particles, the zeta potential should ideally be higher than +20 mV. The charge and zeta potential are just necessary to interact with the alginate/κ-carrageenan long enough for a shell structure to be formed. Therefore, to optimize the particle size, we selected ethanol 15 v/v% as the working concentration.

### 3.2. Optimization of core-shell microparticles (Effect of ratio and concentration)

To optimize the ratio and concentration of each component, we investigated the effect of alginate-κ-carrageenan and phytase concentration on the properties of core-shell microparticles, including encapsulation efficiency and loading capacity. We used a fixed chitosan concentration of 4 w/v% based on previously reported results (Yang et al., 2024). Regarding loading capacity, as the commercial application determines the target, our criteria are 30 w/w% phytase loading capacity. The minimum concentration of alginate-κ-carrageenan to form a shell is 0.15 w/v%, and the loading capacity will decrease if the concentration is too high. So, we selected three conditions, 0.15, 0.30, and 0.45 w/v% of alginate-κ-carrageenan concentrations. Meanwhile, in a previous study (Yang et al., 2024) of phytase-loaded hydrogel beads, controlling the phytase ratio to polysaccharides improved the loading capacity. So, we expected the higher phytase concentration in the phytase-chitosan microparticle to increase the loading capacity. So, we compared three conditions: 4, 6, and 8 w/v% of the phytase concentration in the original phytase-chitosan solutions.

We found that higher alginate-κ-carrageenan concentration increases the encapsulation efficiency, likely due to the formation of a denser outer matrix that reduces phytase leakage during particle formation (**Figure 3(a)**). Also, higher phytase concentrations increase encapsulation efficiency based on a higher driving force for phytase entrapment within the polymer network. Regarding loading capacity, the alginate-κ-carrageenan concentration slightly decreases loading capacity, as the relative proportion of polymer increases compared to phytase. (**Figure 3(b)**). Accordingly, phytase concentration increases the loading capacity because of the slightly higher encapsulation efficiency and phytase ratio. As a result, four samples (phytase 6 and 8 w/v% with alginate/κ-carrageenan 0.30 and 0.45 w/v%) have a higher loading capacity than 30% and higher encapsulation efficiency than 70%. So, we compared those samples to the stability of phytase.

**Figure 3.**
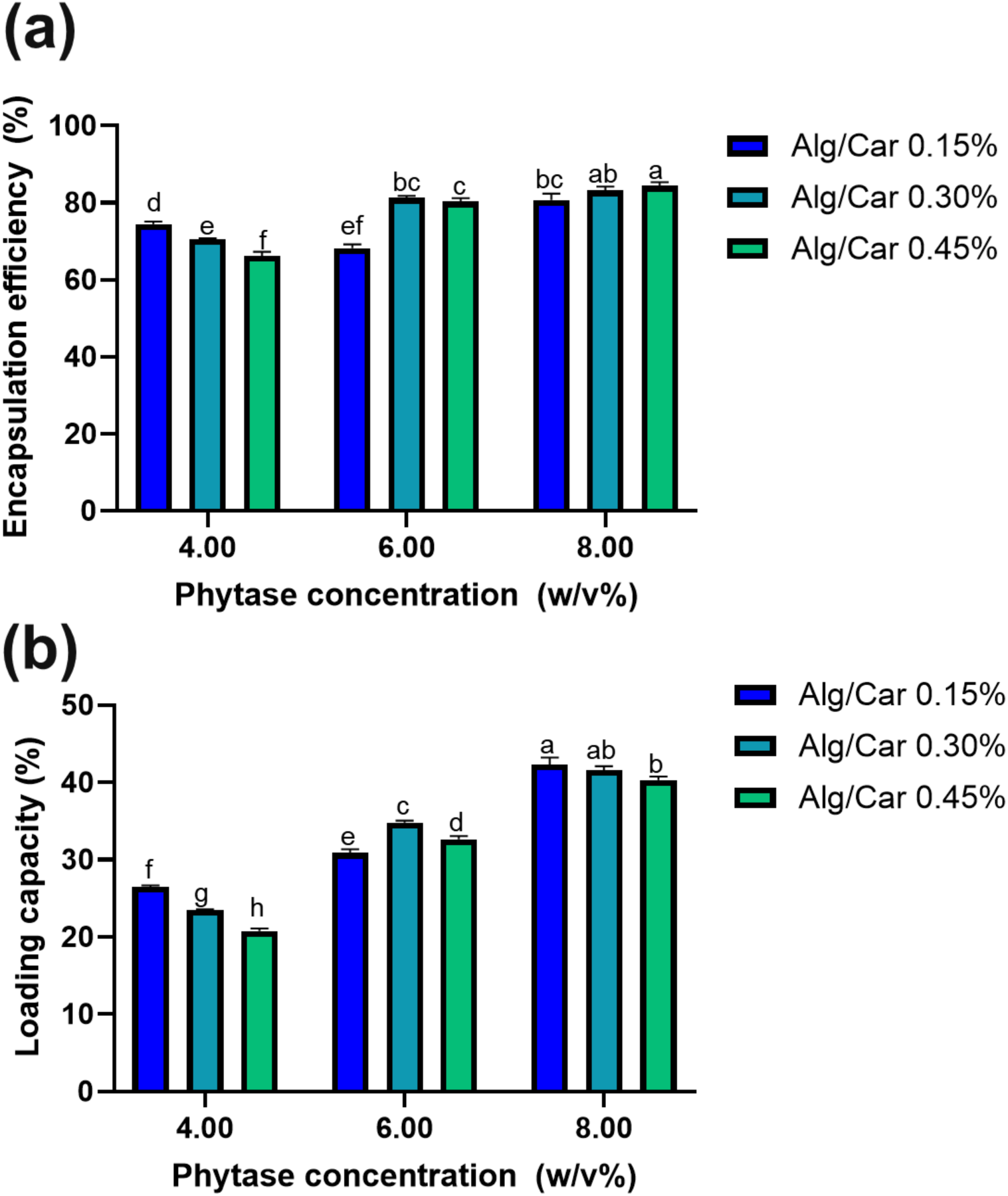
Effect of phytase and alginate/carrageenan concentration on phytase-loaded core-shell microparticles. (a) Encapsulation efficiency and (b) loading capacity of phytase-loaded core-shell microparticles. A connecting letter plot was used at the significance difference level of p ≤ 0.05.

### 3.3. Heat and digestion stability of core-shell microparticles

#### 3.3.1. Effect of heat and acidity on phytase release

Many plant-based foods often require heating, including maize porridge and bouillon, which are heated for at least 5 min at 100 ℃ (S. Wang et al., 2024). Phytase has been found to be temperature sensitive; we anticipated that a core-shell microparticle would protect its activity during heating and at digestive pH levels. To compare the effect of the alginate/κ-carrageenan shell on the stabilization of phytase against high temperature, we measured the release profile after heating at 100 °C for 10 min. We found that creating a core-shell microparticle using a 0.45 w/v% alginate/κ-carrageenan solution produced particles that were better protected from heat-induced activity loss than those made with solutions at 0.15 and 0.30 w/v%. We attribute this to the thicker shell that forms at higher concentration (**Figure 4(a)**).

**Figure 4.**
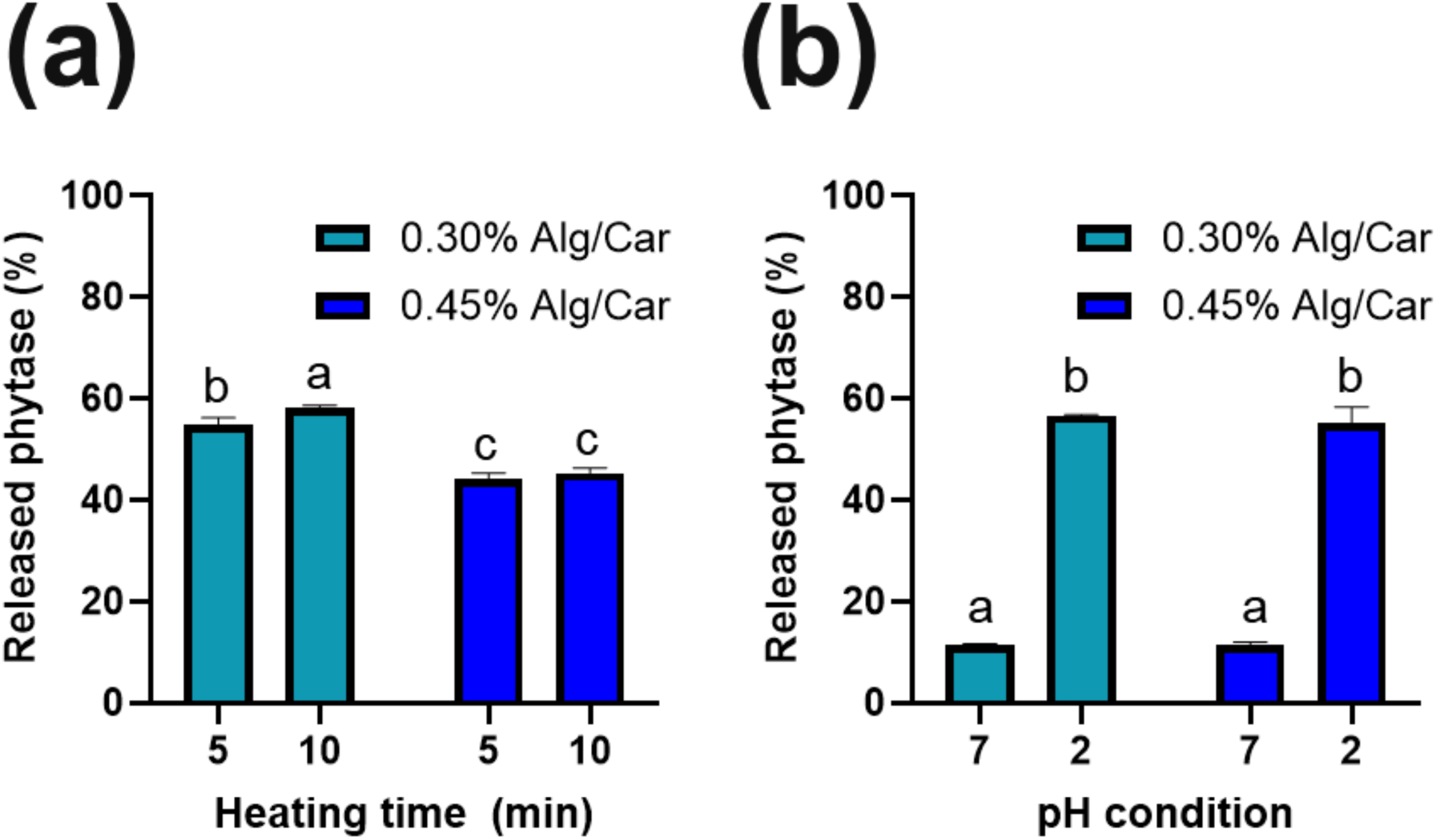
Effect of (a) heat and (b) acidity on the release profile of phytase with respect to the concentration of alginate-carrageenan solution. A connecting letter plot was used at the significance difference level of p ≤ 0.05.

The effect of acidic conditions on enzymes and hydrocolloids includes protein denaturation and active site residue deformation (Lima et al., 2018). Ideally, the release of phytase will not occur until the core-shell microparticles reach the small intestines, we incubated the particles in a pH 2 buffer for 2 h, without pepsin effect. At the same time, we tested pH 7 as a neutral control to compare against acidic exposure. We expected that the thicker shell, which results from higher concentration of alginate and k-carrageenan, would slow down or prevent release; however, the impact was insignificant (p = 0.515 or p > 0.05) (**Figure 4(b)**). At pH 2, alginate and κ-carrageenan have negative charges, and phytase and chitosan have positive charges, resulting in an attractive force between shell and core, while there is a repulsive force between chitosan and phytase. Because we used a lower concentration of shell than core to increase loading capacity, the effect of the repulsive force is stronger than the attractive force. The 0.45 w/v%, because it provided better protection from heat, was selected for further study.

#### 3.3.2. Enzymatic activity of core-shell microparticle after heat and digestion

The enzymatic activity of the optimum formula for the core-shell microparticles (4 w/v% chitosan, 6 w/v% alginate/κ-carrageenan) was tested after heating at 100 °C for 10 min, followed by incubation with gastric fluid, including acidic pH and pepsin, for 2 h. The sample with chitosan 4 w/v%, phytase 6 w/v%, and alginate/κ-carrageenan 0.45 w/v% retained the highest residual activity, which was 71.5 ± 8.4% after heating, this composition for the microparticles also had the highest residual activity after heating and digestion (62.7 ± 4.5%) (**Figure 5**). Based on the residual activity and release profiles, we selected chitosan 4 w/v%, phytase 6 w/v%, alginate/κ-carrageenan 0.45 w/v% as the optimum composition for the microparticle core-shell system.

**Figure 5.**
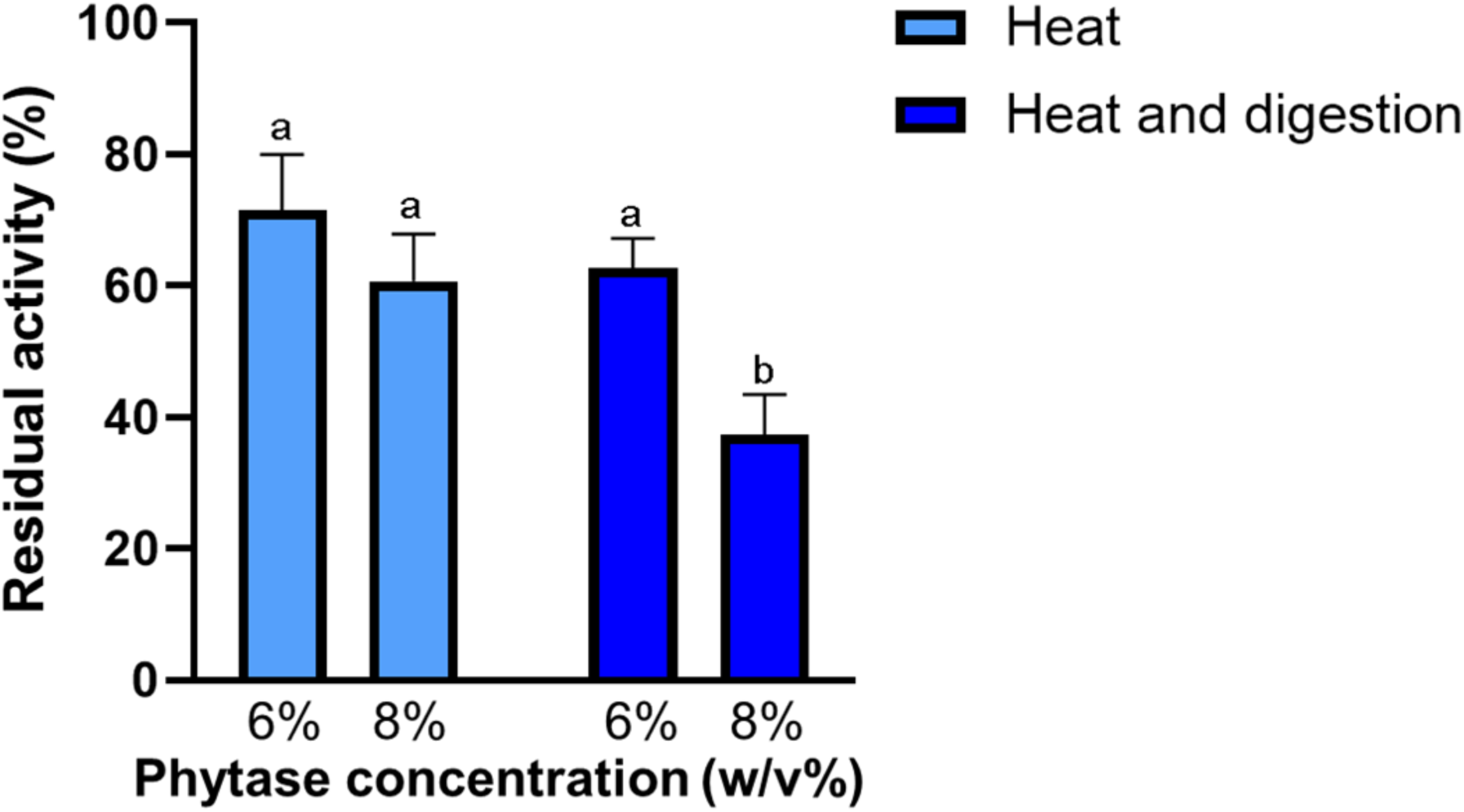
Heat and digestion stability (residual activity) of phytase-loaded core-shell microparticle depending on the phytase concentration after heat for 10 min and gastric digestion incubation for 2 h. A connecting letter plot was used at the significance difference level of p ≤ 0.05.

### 3.4. Characterization of core-shell microparticles

The size of the chitosan-phytate complex increased significantly (p = 0.002) once the alginate-k-carrageenan solution formed a shell around it (**Figure 6(a)**). Using the optimum composition for heat and activity retention after gastric conditions, we prepared the phytase-chitosan/alginate-κ-carrageenan core-shell microparticle and found that the diameter was 3.2 ± 0.3 µm with a uniform distribution (PdI: 0.178 ± 0.118) (**Figure 6(b)**). Compared to the phytase-chitosan complex, the diameter increased by 1.8-fold upon formation of phytase-chitosan microparticles. Further complexation into core-shell microparticles resulted in an additional size increase, making particles that were 2.7 times larger than the phytase-chitosan microparticles. We confirmed the size of the final product through SEM analysis and measured the average diameter using ImageJ. The SEM image showed uniform and puckered particles with a microsize (3.15 ± 0.70 µm) (**Figure 6(c)**). The puckering is an artifact of the spray-drying process, which makes the core-shell microparticles more compact by removing any water molecules between phytase-chitosan microparticles and the shells.

**Figure 6.**
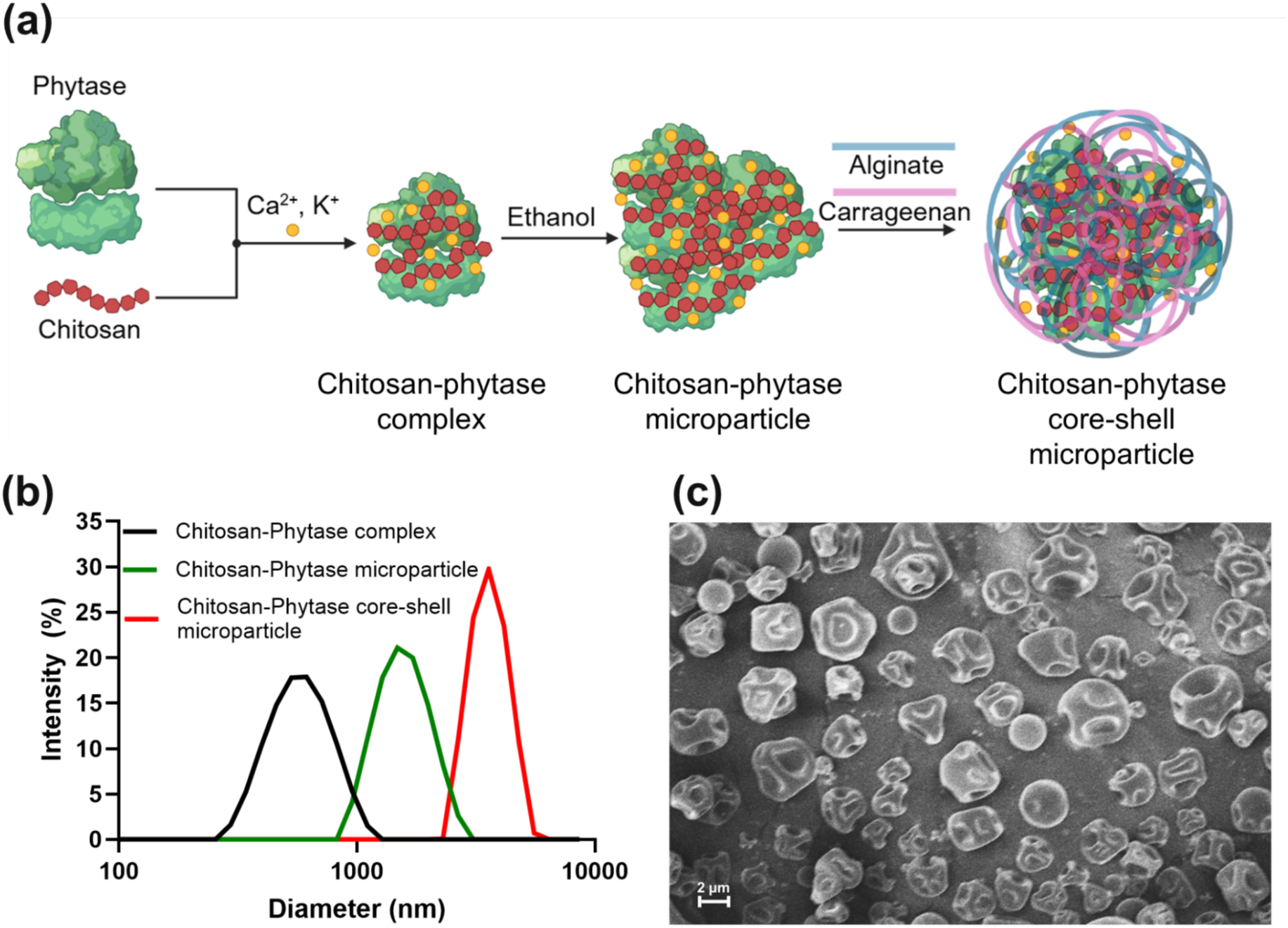
(a) Graphical illustration of preparation steps of chitosan-phytase core-shell microparticles. (b) Size distribution of chitosan-phytase complex, chitosan-phytase microparticle, and chitosan-phytase core-shell microparticle. (c) SEM image of phytase-loaded core-shell microparticles (scale bar = 2 µm).

To better understand the nature of the interactions of the materials within the core-shell microparticle, we compared the infrared spectra of alginate, κ-carrageenan, phytase, and chitosan (**Figure 7(a)**). Compared to the phytase, the peak intensity at 1404 cm^-1^ decreased, and the peak at 1030 cm^—1^ decreased and shifted to 1065 cm^-1^ (**Figure 7(b)**). The absence of new peaks associated with the polysaccharides and the phytase in the loaded particle suggested that the proteins and polysaccharides are attracted primarily through electrostatic interactions and not new covalent bonds. This shift and intensity decrease is most likely associated with hydrogen bonds, hydrophobic forces, and electrostatic interactions.

**Figure 7.**
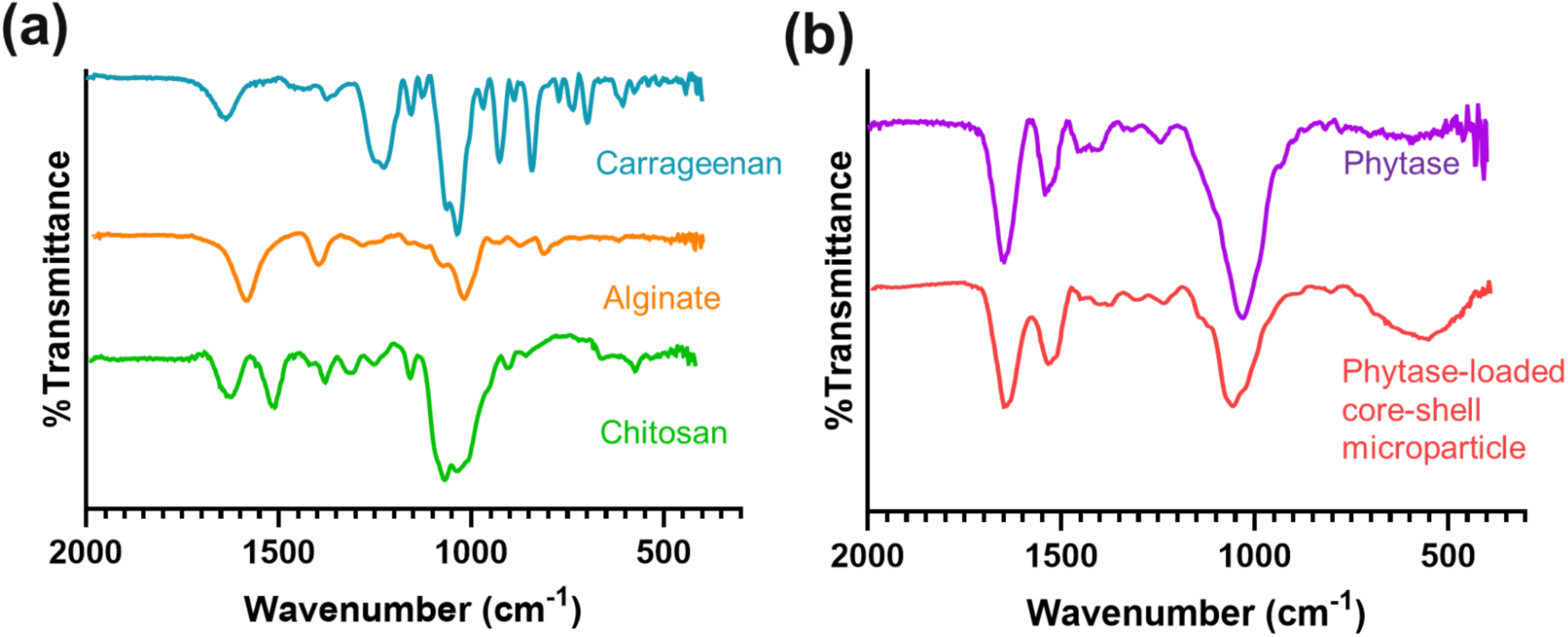
FT-IR result of (a) the polysaccharides, including carrageenan, alginate, and chitosan, and (b) phytase and phytase-loaded core-shell microparticles.

### 3.5. Enzymatic activity of core-shell microparticles in food matrix

Based on the effect of heat and digestion on the enzyme integrity, we used our optimized core-shell microparticles for phytate hydrolysis in a food matrix (**Figure 8(a)**). Ideally, phytase should degrade phytate before reaching the small intestine, releasing the minerals it has complexed at the location where they are absorbed; therefore, phytate hydrolysis will be evaluated during food processing and during gastric digestion. Furthermore, to determine the amount of phytase-loaded core-shell microparticle powder for food application, specifically maize porridge, we had to calculate the phytase activity unit (FTU)of our sample. The Food and Agriculture Organization (FAO) has defined 1 FTU as the amount of phytase that releases 1 μmol of inorganic phosphorus per minute from 5 mM sodium phytate at 37 °C and pH 5.5. In a food matrix, 100 g of whole maize powder has ∼0.6 g phytate, and maize porridge includes 30∼40 w/v% maize powder. Under these conditions, the final phytate concentration in maize porridge was 1.8 mM, which allowed us to redefine 1 FTU as the amount of phytase that releases 1 μmol of inorganic phosphorus per minute from 1.8 mM phytate at 37 °C and under gastric conditions, including pepsin.

**Figure 8.**
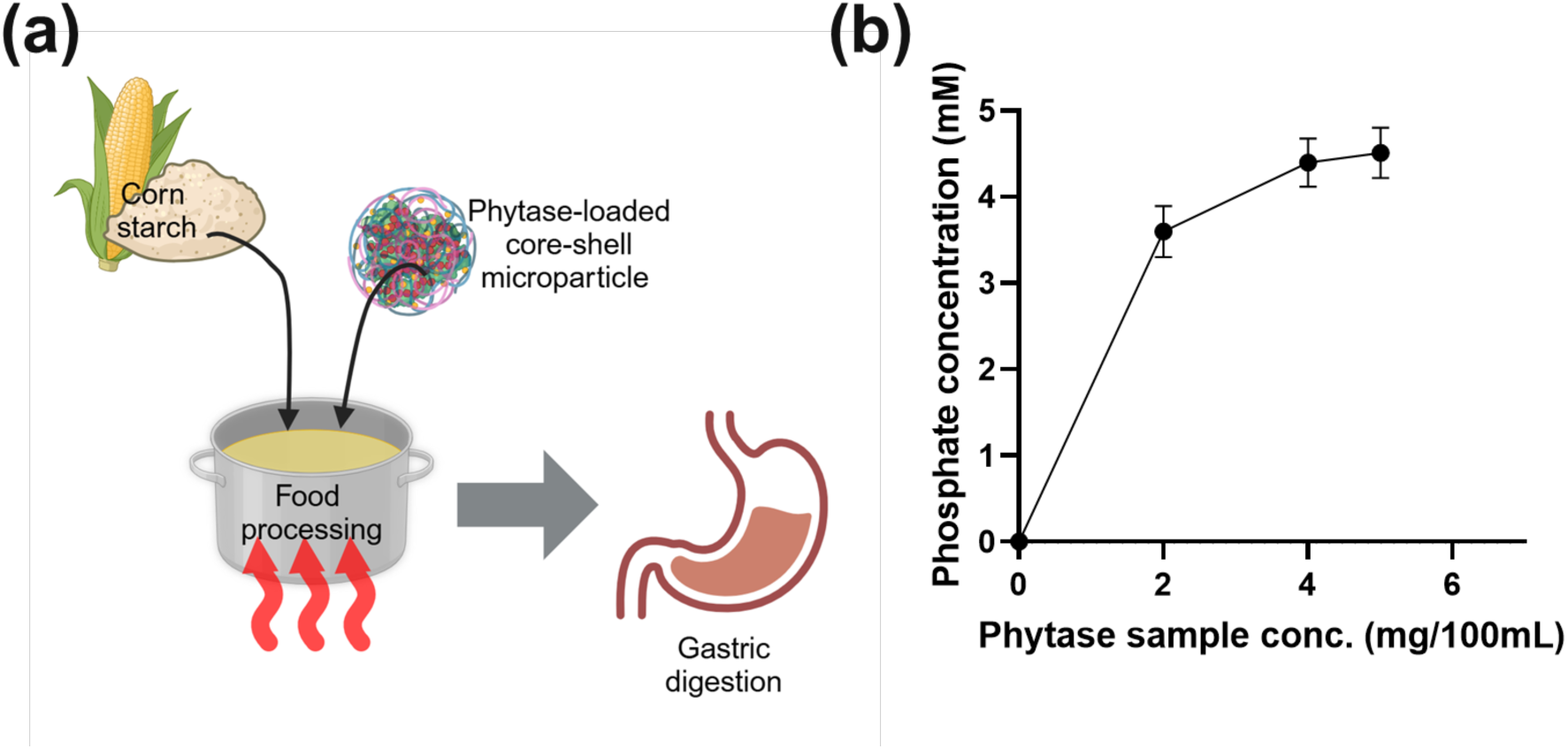
(a) Graphical illustration of food processing and digestion. (b) The concentration of phosphate produced by phytate hydrolysis depends on the loaded phytase powder concentration.

Theoretically, 1 mole of phytate (IP6) can release 6 moles of inorganic phosphate upon complete hydrolysis by phytase (IP6 → IP5 → IP4 → … → IP0). However, most phytases are optimized to degrade phytate to intermediate products (e.g., IP4, IP3), liberating a significant portion of phosphate but not all. Therefore, if phytase hydrolyzes IP6 to IP5: phosphate = 1.8 mM. If phytase hydrolyzes IP6 to IP4: phosphate = 3.6 mM. If phytase hydrolyzes IP6 to IP0: phosphate = 10.8 mM. In our experiments, phosphate concentration plateaued at 4.5 mM (**Figure 8(b)**), indicating that under these conditions, hydrolysis occurred to IP4 or IP3 due to enzyme specificity (3-phytase) and potential enzyme inhibition. We calculated that 6 FTU of phytase is required to degrade 1.8 mM phytate to IP4, producing 3.6 mM phosphate. Each milligram of our phytase sample has an activity of 3 FTU; therefore, 2 mg of the core-shell microparticles is required for a single maize meal to achieve maximum phytate degradation by phytase.

## 4. Conclusions

A core-shell microparticle was fabricated to maintain the catalytic activity of phytase while protecting it from high processing temperatures, acidity, and digestion conditions. We optimized the fabrication conditions for the phytase-chitosan core-shell microparticle. The optimized core-shell microparticle has a loading capacity of 32.7% with a high encapsulation efficiency of 80.3% and a uniform microparticle size of 3.2 ± 0.3 µm and a narrow polydispersity of 0.178 ± 0.118. The resulting core-shell microparticle showed good tolerance against heat and acidic conditions. Loaded phytase retained 62.7% of its enzymatic activity even after heat treatment and digestion conditions. Finally, we found that only 2 mg of our sample can hydrolyze phytate in a single maize meal to help mineral absorption. These results suggest that the core-shell microparticles are suitable for retaining enzyme activity against various stressors, making it an advantageous method for food fortification, such as in maize porridge.

## Author CRediT statement

**Eunhye Yang:** Conceptualization, investigation, formal analysis, visualization, writing-original draft, reviewing and editing. **Waritsara Khongkomolsakul:** Conceptualization, investigation, writing review, and editing. **Younas Dadmohammadi:** Project conceptualization, supervision, funding acquisition, writing review, and editing. **Alireza Abbaspourrad:** Project conceptualization and administration, supervision, funding acquisition, writing review, and editing

## Generative Artificial Intelligence (AI) Statement

The authors certify that generative AI was not used in preparing this article. Non-generative AI, such as spelling and grammar checkers in Office 365 and Google Docs, and citation managing software, was used. All instances when non-generative AI was used were reviewed by the authors and editors.

## Acknowledgments

This work was carried out with the support of the Nutrition R&D Program (INV-043930) provided by the Bill & Melinda Gates Foundation (WA, US).

